# Sex-dependent effects of chronic exercise on cognitive flexibility in aging mice

**DOI:** 10.1101/2020.06.10.145136

**Authors:** Annabel K. Short, Viet Bui, Isabel C. Zbukvic, Anthony J. Hannan, Terence Y. Pang, Jee Hyun Kim

**Affiliations:** Florey Department of Neuroscience and Mental Health University of Melbourne, Parkville, VIC 3052 Australia; Mental Health theme, The Florey Institute of Neuroscience and Mental Health, Parkville, VIC 3052 Australia

**Keywords:** sex characteristics, cognition, exercise, hippocampus, *Bdnf*, reinstatement

## Abstract

Cognitive impairments associated with advanced age are a growing concern in our aging society. Such impairments are associated with alterations in brain structure and function, especially in the hippocampus, which changes to experience throughout life. It is well-known that regular exercise can maintain hippocampus volume. The hippocampus is critical for cognitive flexibility involved with extinction and reinstatement of conditioned fear. Therefore, we asked whether voluntary chronic exercise in middle-aged mice can improve extinction and/or reinstatement of conditioned fear compared to standard housing. Eight-month-old male and female C57Bl/6J mice had access to a running wheel or remained in standard housing until 11 months of age. Alongside control standard-housed young adult (3-month-old) mice, they received tone-footshock pairings, which were subsequently extinguished with tone-alone presentations the next day. Half of the mice then received a reminder treatment in the form of a single footshock. Both male and female 11-month-old mice housed in standard conditions exhibited impaired reinstatement compared to young adult mice. However, for males that had access to a running wheel from 8 months of age, the reminder treatment rescued reinstatement ability. This was not observed in females. Additionally, exercise during middle age in both sexes increased expression of *Bdnf* mRNA in the hippocampus, specifically exon 4 mRNA. These results show that, at least for males, physical exercise is beneficial for reducing age-related decline in cognitive abilities. Despite not rescuing their impaired reinstatement, exercise also increased *Bdnf* gene expression in the female hippocampus, which could potentially benefit other forms of hippocampal-dependent cognition.

## 1. Introduction

Aging is associated with reduced executive functions, with cognitive flexibility being one of the most impaired facets of intelligence due to age (Salthouse, 1996). Cognitive flexibility is the ability to adapt to a changing environment and is often tested using reversal or set shifting tasks (Dajani & Uddin, 2015; Scott, 1962). Currently, medical interventions for cognitive impairments associated with aging are limited. Thus, identifying new ways to delay age-associated cognitive problems is vital in our aging society.

People with a history of a physically active lifestyle are known to have some resilience to the effects of normal aging on cognition. In humans, the correlation between aerobic exercise and improved cognition is strong and exercise appears to be effective across the lifespan (Chaddock et al., 2010; Erickson et al., 2011; Herting & Nagel, 2012; Kleemeyer et al., 2016; Rosano et al., 2017; Stillman et al., 2018; Thomas et al., 2016). This has significant implications for older adults who face heightened risk of dementia (Erickson et al., 2011; Kleemeyer et al., 2016; Rosano et al., 2017).

The effects of exercise on cognition has been linked with hippocampal function (Rubin, Watson, Duff, & Cohen, 2014), which is important for spatial tasks and cognitive flexibility (Burghardt, Park, Hen, & Fenton, 2012). Although many studies highlight changes in the hippocampus volume and connectivity as the neural correlate for exercise effects in humans (Chaddock et al., 2010; Erickson et al., 2011; Herting & Nagel, 2012; Kleemeyer et al., 2016; Rosano et al., 2017), the molecular correlates are poorly understood. Rodent models have been useful in this regard and have provided additional insights. For example, chronic exercise can alleviate the decrease of hippocampal neurogenesis and synaptic plasticity in aging rodents (Anacker & Hen, 2017; van Praag, Shubert, Zhao, & Gage, 2005).

Importantly, the hippocampus is a sexually-dimorphic brain region in rodents and humans, dependent on hormonal cycle stage (reviewed in Yagi & Galea, 2019). Indeed, sex differences in hippocampus-dependent learning have been widely reported in rodents and humans, with males typically showing superior spatial learning (Jonasson, 2005; Linn & Petersen, 1985; Voyer, Voyer, & Bryden, 1995). However, whether there are sex differences in cognitive decline with age is unclear in humans (Ferreira, Ferreira Santos-Galduróz, Ferri, & Fernandes Galduróz, 2014; Karlsson, Thorvaldsson, Skoog, Gudmundsson, & Johansson, 2015; McCarrey, An, Kitner-Triolo, Ferrucci, & Resnick, 2016; Workman, Healey, Carlotto, & Lacreuse, 2019; Zaninotto, Batty, Allerhand, & Deary, 2018). In rodents, there are very few reports of sex differences in non-pathological cognitive decline with age (Zanos et al., 2015). Sex differences are observed following exercise in hippocampus-dependent tasks, although rodent and human findings differ. A meta-analysis in humans reported sex-differences in the level of cognitive improvements following exercise, with females having greater improvements especially after aerobic training (Barha, Davis, Falck, Nagamatsu, & Liu-Ambrose, 2017). A similar meta-analysis in rodents describes no sex differences in spatial tasks following aerobic training, but greater improvements in non-spatial cognitive tasks in males (Barha, Falck, Davis, Nagamatsu, & Liu-Ambrose, 2017). Taken together, it is clear that sex-specific effects on cognition and underlying neurobiology need further examination.

The aim of the present study was to examine the impact of exercise on cognitive flexibility and hippocampus in aging male and female mice. Cognitive flexibility was assessed using reinstatement following extinction of conditioned fear. Mice were first conditioned with a tone conditioned stimulus (CS) that was paired with a footshock unconditioned stimulus (US), which led to freezing to the CS as a measure of emotional memory of the conditioning session (CS-US). Then the CS was presented repeatedly without the US, which decreases the freezing to the CS to form the extinction memory (CS-no US). When tested in the same context as extinction, the CS-no US memory is typically retrieved, evidenced by low levels of freezing. However, a single reminder footshock can facilitate the retrieval of the conditioning memory and lead to high freezing (i.e., reinstatement). Taken together, reinstatement can test cognitive flexibility because it requires flexible retrieval of the extinction vs conditioning memory depending on the reminder given (Anacker & Hen, 2017; Kim & Richardson, 2010; Short et al., 2016). While it is not the typical model to study cognitive flexibility, it is widely agreed that expression of reinstatement requires a complex understanding of environmental cues to be flexible in the choices of responses, and deficit in such cognitive flexibility may be related to persistence of fear observed in anxiety disorders (Bouton, 2002; Ganella & Kim, 2014; Giovanello, Schnyer, & Verfaellie, 2009; Kim & Richardson, 2010; Maren, Phan, & Liberzon, 2013). Consistent with these ideas, hippocampal lesions impair reinstatement (Frohardt, Guarraci, & Bouton, 2000; Wilson, Brooks, & Bouton, 1995). In those studies, it was not the context-specificity of reinstatement that was abolished, but reinstatement itself was abolished in the same context as where the reminder is given (Frohardt, Guarraci, & Bouton, 2000; Wilson, Brooks, & Bouton, 1995). Those results demonstrate that similar to cognitive flexibility, the hippocampus is a critical structure also for reinstatement. This model is particularly appropriate for this study because previous studies have found that reinstatement is both sex- and age-specific in rodents (Kim & Richardson, 2007; Matsuda et al., 2015; Park, Ganella, & Kim, 2017a; Voulo & Parsons, 2017), highlighting that it is sensitive to age and sex effects.

In addition, we examined hippocampal brain-derived neurotrophic factor (*Bdnf)* gene expression of these mice to see if the beneficial effects of exercise are reflected at a molecular level. Increased hippocampal *Bdnf* expression in freely exercising rodents is a well-established molecular correlate for the exercise-associated benefits on brain and behavior (Berchtold, Chinn, Chou, Kesslak, & Cotman, 2005; Berchtold, Kesslak, Pike, Adlard, & Cotman, 2001; Cotman, Berchtold, & Christie, 2007; Neeper, Gómez-Pinilla, Choi, & Cotman, 1996). In particular, *Bdnf* exon 4 transcript expression has been implicated in extinction of conditioned fear in rodents (Baker-Andresen, Flavell, Li, & Bredy, 2013; Bredy et al., 2007).

We chose mice at 3 months of age as young adult baseline controls, which is a well-established age of young adulthood based on hormone and cognitive development (Bell, 2018; Madsen & Kim, 2016). In addition, we showed previously that 3-month-old mice reliably display reinstatement of extinguished fear (Chen et al., 2016). For the aged group, we chose 11-month-olds to test reinstatement at an age with documented cognitive decline in mice (Lynch, Rex, & Gall, 2006; Marlatt, Potter, Lucassen, & van Praag, 2012), while avoiding the onset of reproductive senescence in female mice occurring at 12 months of age (Diaz Brinton, 2012; Koebele & Bimonte-Nelson, 2016) to ensure potential sex differences during aging can be observed. We hypothesized that an age-related impairment of reinstatement and hippocampal BDNF gene expression would be reversed by voluntary exercise during middle age. Based on meta-analysis in mice (Bahar, Falck et al., 2017), we also anticipated that exercise effects would be greater in males than females.

## 2. Material and Methods

### 2.1. Animals

C57Bl/6J mice at 8 weeks of age were purchased from the Animal Resources Centre (Murdoch, WA, Australia) in 3 cohorts. They were aged to 8 months, and assigned to either Standard Housing or Exercise. All animals were group housed (2-3 mice, males and females separately) in cages with a floor area of 1862 cm^2^ (GR1800, Techniplast, Australia). Exercising animals had access to 1 running wheel per mouse for 3 months until behavioral testing commenced at 11 months of age. Exercise wheels were removed the day before behavioral testing. C57Bl/6J mice also purchased at 8 weeks of age from ARC served as Young controls. They were housed similarly to Standard Housing aging mice with no access to running wheels and were tested at 3 months of age concurrently with the aged mice. All cages were lined with sawdust and two tissues provided for nesting with food and water *ad libitum*. Temperature and humidity were controlled at 22°C and 45%, respectively. Mice were maintained on a 12-h light/dark cycle (lights on at 0700 hours) and bedding changed weekly. All experiments were approved and performed in accordance with the guidelines of the Florey Institute of Neuroscience and Mental Health Animal Ethics Committee.

### 2.2. Behavior

#### 2.2.1. Apparatus

Fear conditioning chambers were rectangular (31.8 cm × 25.4 cm × 26.7 cm) with stainless steel grid floors with 36 rods (3.2 x 7.9mm), equipped with a Med Associates VideoFreeze system (Med Associates, USA). A constant-current shock generator was used to deliver electric shock (0.7mA, 1s) (unconditioned stimulus, US) to the floor of the chambers as required. A programmable tone generator, speaker and sound calibration package was used to deliver auditory tone (volume: 80 dB; frequency: 5000 Hz) (conditioned stimulus, CS). In order to create two different contexts the chambers differed in appearance as described previously (Handford et al. 2014). One type of context contained a white house light, curved striped walls and a tray lined with paper towels placed 10 cm beneath the grid floor. The second context had no house light, and contained no wall insert and a tray lined with clean bedding beneath the grid floor. Individual chambers were enclosed in sound-attenuating boxes within the same room. The experimental room in which the chambers were located was brightly lit with overhead lights. Animals were randomly assigned to different starting contexts.

#### 2.2.2. Conditioning

Mice at either 3 or 11 months-of-age concurrently received fear conditioning. Mice were placed in the chamber, and baseline freezing was measured for 2 minutes. Following this, all animals received 6 tone–footshock (CS-US) pairings. Each pairing consisted of a 10s tone that co-terminated with a 1s shock (0.7mA). Inter-trial intervals (ITIs) ranged from 85 to 135s, with an average of 110s. Two minutes following the last presentation, the mouse was removed from the chamber and placed back into the home cage.

#### 2.2.3. Extinction

The day following conditioning, mice were tested for their fear CS memory by being placed in a different context to that in which they were conditioned. Mice were allowed a 2 min period during which baseline freezing was measured. They then received 45 presentations of a 10s tone in the absence of the shock with ITI of 10s.

#### 2.2.4. Reinstatement and test

The day following extinction, the mice were divided into two groups. One group (Reminder) received a single reminder shock (0.7mA, 1s) in the extinction context, and the other group (No Reminder) was placed in the extinction context but did not receive any shock. The following day the rats were tested as described for extinction (2 min baseline followed by 45 presentations of a 10s tone in the absence of the shock with ITI of 10s (i.e. extinction) in the same context as where extinction and reminder occurred.

### 2.3. Gene expression analysis

#### 2.3.1. RNA extraction

One-week following the reinstatement test, whole hippocampi were micro-dissected from animals from the second cohort of males (3-months standard n=4; 11-months standard n=3; 11-months running n=3) and females (3-months standard n=4; 11-months standard n=3; 11-months running n=3). RNA was extracted using QIAGEN RNeasy Mini extraction kit as per manufacturer’s instructions (QIAGEN, VIC, Australia). Tissue samples were disrupted using a Diagenode Biorupter (UCD-300; Life Research, VIC, Australia) in the QIAGEN lysis buffer. On-column DNAse1 treatment was performed, and RNA was eluted in 50μL of RNase-free water. RNA concentrations and purity were determined using Nanodrop spectrograph (2000c Thermo Scientific, Wilmington, DE, USA). Samples were then stored at −80°C until required.

#### 2.3.2. Reverse transcription

Thawed RNA (1000ng) was reverse transcribed using Superscript VILO cDNA synthesis kit (Invitrogen, Life Technologies, VIC, Australia). Reverse transcription PCR was performed in a thermal cycler (Takara Shuzo, Tokyo, Japan) using 1 x cycle of the following programme: 25°C for 10 minutes, 48°C for 30 minutes and 95°C for 5 minutes. Samples were then stored at −20 °C.

#### 2.3.3. Real-time quantitative PCR (qPCR)

Levels of gene expression in the tissue was quantified using qPCR using either the Viia 7 Real-Time PCR system (Applied Biosystems, CA, USA). Reactions were made using: SYBR green, 10μL (S4438, Sigma-Aldrich, NSW, Australia), ROX reference dye 0.2 μL (12223-012, Invitrogen, VIC, Australia), Forward and reverse primers (20μM) 0.5-1.5μL each, cDNA 50ng in 5μL, DNAse-free H_2_O up to 20μL. Cyclophillin was used as the endogenous control. Primer sequences:

Cyclophillin forward 5’CCCACCGTGTTCTTCGACA3’

Reverse 5’CCAGTGCTCAGAGCTCGAAA3’.

*Bdnf* total forward 5’ GCGCCCATGAAAGAAGTAAA 3’

Reverse 5’ TCGTCAGACCTCTCGAACCT 3’

*Bdnf* Exon 4 forward 5’ CAGAGCAGCTGCCTTGATGTT 3’

Reverse 5’ GCCTTGTCCGTGGACGTTTA 3’

Primers were ordered from Sigma-Aldrich (Castle Hill, NSW). Optimal primer dilutions and amplification efficiencies had previously been determined by TYP and AKS. PCRs were run on the following programme: 50°C for 2 minutes, 95°C for 10 minutes, followed by 40x cycles of 95°C for 15 seconds and 60°C for 1 minute. The expression levels of the target genes were determined using comparative Ct (ΔΔCt) method and normalized to the mean expression of the young adult male control group.

### 2.4. Statistical analysis

#### 2.4.1. Overall

Statistics were computed using SPSS statistics version 22.0 (IBM, Armonk NY, USA) and GraphPad prism 6.0 (GraphPad software, Inc., La Jolla, CA. All behaviors were analyzed as described previously (Madsen, Guerin, & Kim, 2017; Short et al., 2017). For conditioning, percentage freezing during the first 9 seconds of each CS-US trial was analyzed to avoid analyzing shock-related movement. For extinction, percentage freezing reported is based on 10s of tone blocked into average freezing of 5 tones to produce 9 blocks. For test, percentage freezing reported is based on 10s of tone presentations averaged across the whole test. Three-way repeated measures analyses of variance (ANOVA) was employed to analyze potential effects of Sex, Treatment, Reminder and interactions between these factors on freezing levels. Three-way ANOVA was employed to analyze potential effects of Sex, Treatment, Reminder and interactions on the levels of mRNA expression. Main effects were followed up with Tukey’s multiple comparisons *post hoc t*ests. Where there were significant interactions *post hoc* Bonferroni t-tests were performed.

#### 2.4.2. Baseline freezing

At conditioning, there was a significant effect of Sex on baseline freezing (F_(1,72)_=4.05, p<0.05) with males freezing more than females (Figure 1a). There were no other effects or interactions (p > 0.05). At extinction (Figure 1b), there were no effects or interactions (p >0.05). At test, there was an effect of Reminder on baseline freezing (F_(1,72)_=28.38, p<0.001), with mice that received the reminder footshock freezing more than those that did not (Figure 1c). There were no other effects or interactions (p >0.05). To control for pre-existing differences in baseline freezing, CS-elicited freezing for each behavioral session was analyzed using analyses of co-variance (ANCOVA) with baseline freezing levels as a co-variate as described in previous studies (Kim & Richardson, 2007; Park et al., 2017a). However, the results of ANCOVA did not differ from results of ANOVA (i.e., without baseline as a co-variate), therefore, we report the results of ANOVA below.

**Figure 1.**
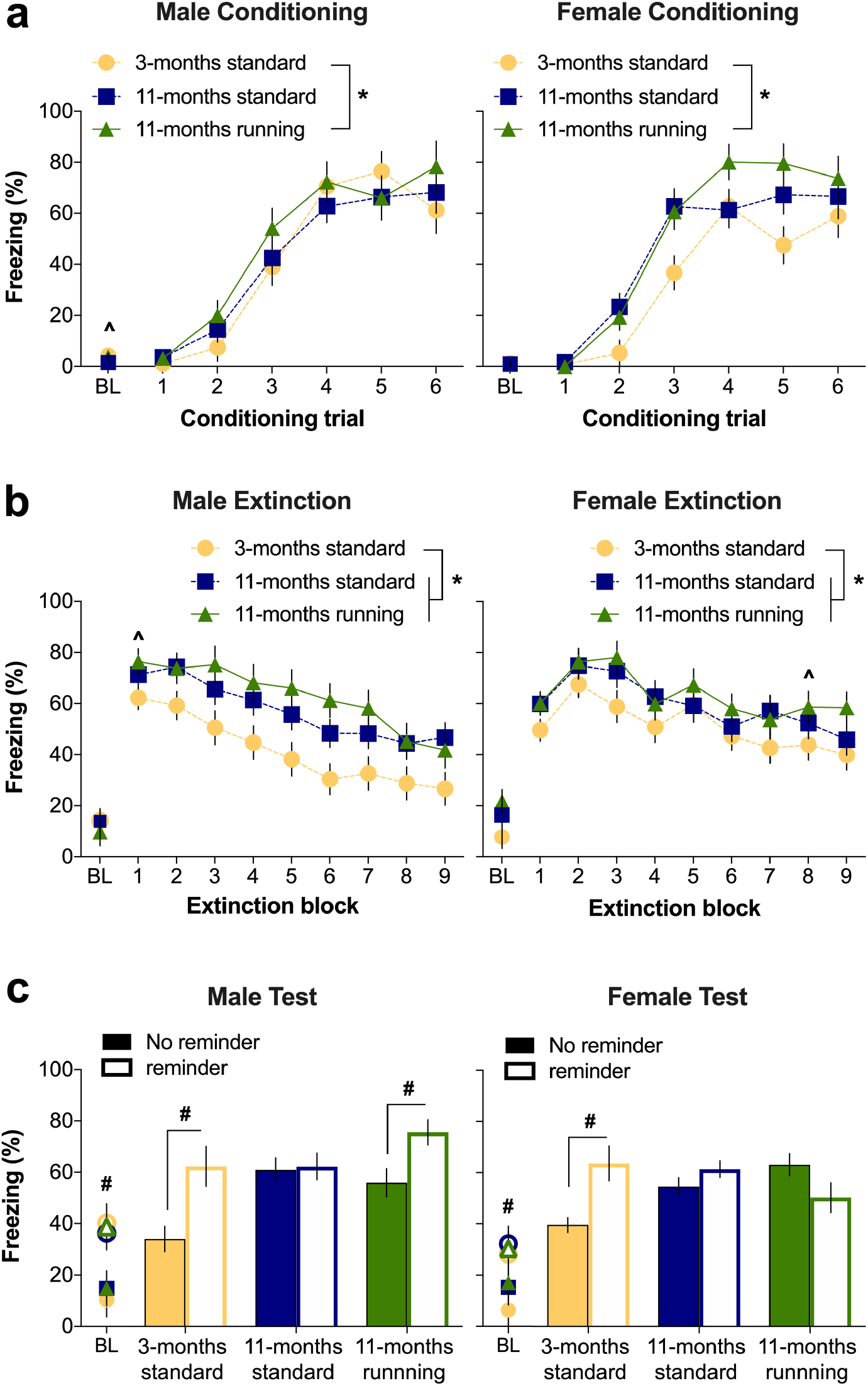
Effect of age and exercise on fear conditioning, extinction, and reinstatement test (Mean ± SEM). (a) While all mice increased their CS-elicited freezing across conditioning trials, aged mice with access to running wheels freeze more when compared to young mice regardless of sex (*p<0.05 main effect of Treatment with post-hoc Tukey comparisons; ^p<0.05 main effect of Sex). (b) Aging elevates CS-elicited freezing levels during extinction in males and females (*p<0.05 main effect of Treatment with significant post-hoc differences). In addition, males overall display greater fear retrieval in the beginning of extinction followed by a steeper extinction curve (i.e., accelerated extinction learning) compared to females, however, there is no sex difference by the final extinction block 9 (^p<0.05 Sex X Block interaction with significant post-hoc effect of Sex). (c) In males, reinstatement is impaired in aged animals living in standard housing, which is rescued by access to running wheels. In females, reinstatement is impaired in aged mice regardless of access to running wheels (#p<0.05 Treatment X Sex X Reminder interaction with significant post-hoc effect of Reminder or main effect of Reminder). Conditioning and extinction: males; n=12-16 and females; n=14-15 per group. Reinstatement test: males; n=4-9 and females; n=7-8 per group.

## 3. Results

### 3.1. Aging impairs cognitive flexibility that is rescued with exercise only in males

Repeated measures (RM) ANOVA of CS-elicited freezing during conditioning (Figure 1a) revealed main effects of Trial (F_(5,360)_=147.99, p<0.001) and Treatment (F_(2,72)_=4.26, p<0.05). Tukey *post hoc* tests revealed that aged mice with access to running wheels froze more than young mice (p<0.01). There were no effects of Sex, Reminder or interactions between any of the factors (p>0.05). These results indicate that while all mice increased their CS-elicited freezing across conditioning trials, aged mice with access to running wheels freeze more when compared to young mice regardless of Sex.

Three-way RM ANOVA of CS-elicited freezing during extinction (Figure 1b) revealed main effects of trial Block (F_(8,576)_=35.91, p<0.001) and Treatment (F_(2,72)_=6.70, p=0.002). Tukey *post hoc* tests revealed that both groups of aged mice froze more than young mice (p<0.05). There was a Block x Sex interaction (F_(8,576)_=5.13, p<0.001) with *post hoc* tests showing males freezing more than females at block 1 (p<0.001), then females freezing more than males at block 8 (p<0.05) but no other blocks showing sex effects (ps>0.05). There were no other effects or interactions (ps>0.05). These results suggest that regardless of Exercise or Sex, aging elevates CS-elicited freezing levels during extinction. In addition, males overall display greater fear retrieval in the beginning of extinction followed by a steeper extinction curve (i.e., accelerated extinction learning) compared to females. However, there is no sex difference by the final extinction block 9.

Three-way ANOVA of CS-elicited freezing during test (Figure 1c) showed main effects of Reminder (F_(1,72)_=13.28, p<0.005) and Treatment (F_(2,72)_=5.41, p<0.01), with Tukey post hoc tests showing that young mice froze less than aged mice (p<0.05). There also were Treatment x Reminder interaction (F_(2,72)_=6.01, p<0.005), and Treatment x Reminder x Sex interaction (F_(2,72)_=3.29, p<0.05) but no other interactions (p>0.05). To understand the 3-way interaction, the effect of Reminder was tested via Bonferroni-corrected t-tests per Sex and per Treatment. In males, the reminder effectively reinstated extinguished fear in young (p<0.01) and running wheel housed aged mice (p=0.05) mice but not in standard housed aged mice (p>0.05). In females, only young animals showed reinstatement (p<0.01), with no Reminder effect in both aged groups (p>0.05). Taken together, this suggests that aging decreases cognitive flexibility in both sexes, and that this can be rescued with exercise in males but not in females.

### 3.2. Exercise up-regulates hippocampal Bdnf mRNA levels in male and female mice

One-week following the reinstatement test, whole hippocampi were micro-dissected from animals from the second cohort of males. While there was no significant Sex x Treatment interaction (F_(2,14)_=1.945, p=0.17) nor effect of Sex (F_(1,14)_=3.77, p=0.07) for total *Bdnf* (Figure 2a), there was an overall effect of Treatment (F_(2,14)_=6.38, p=0.01). Tukey’s *post hoc* comparisons revealed that aged running mice had higher levels of total *Bdnf* when compared to aged standard (p<0.01). There was also a strong trend for its heightened levels in aged running mice compared to young standard (p=0.056). Similarly, for *Bdnf* exon 4 (Figure 2b) there was no Sex x Treatment interaction (F_(2,14)_=0.863, p =0.443) nor effect of Sex (F_(1,14)_=1.77, p =0.21), but there was a significant effect of Treatment (F_(2,14)_=5.67, p =0.01). Tukey’s post hoc comparisons revealed increased *Bdnf* exon 4 mRNA expression in aged running animals compared to young standard and aged standard housed animals, separately (p<0.05).

**Figure 2.**
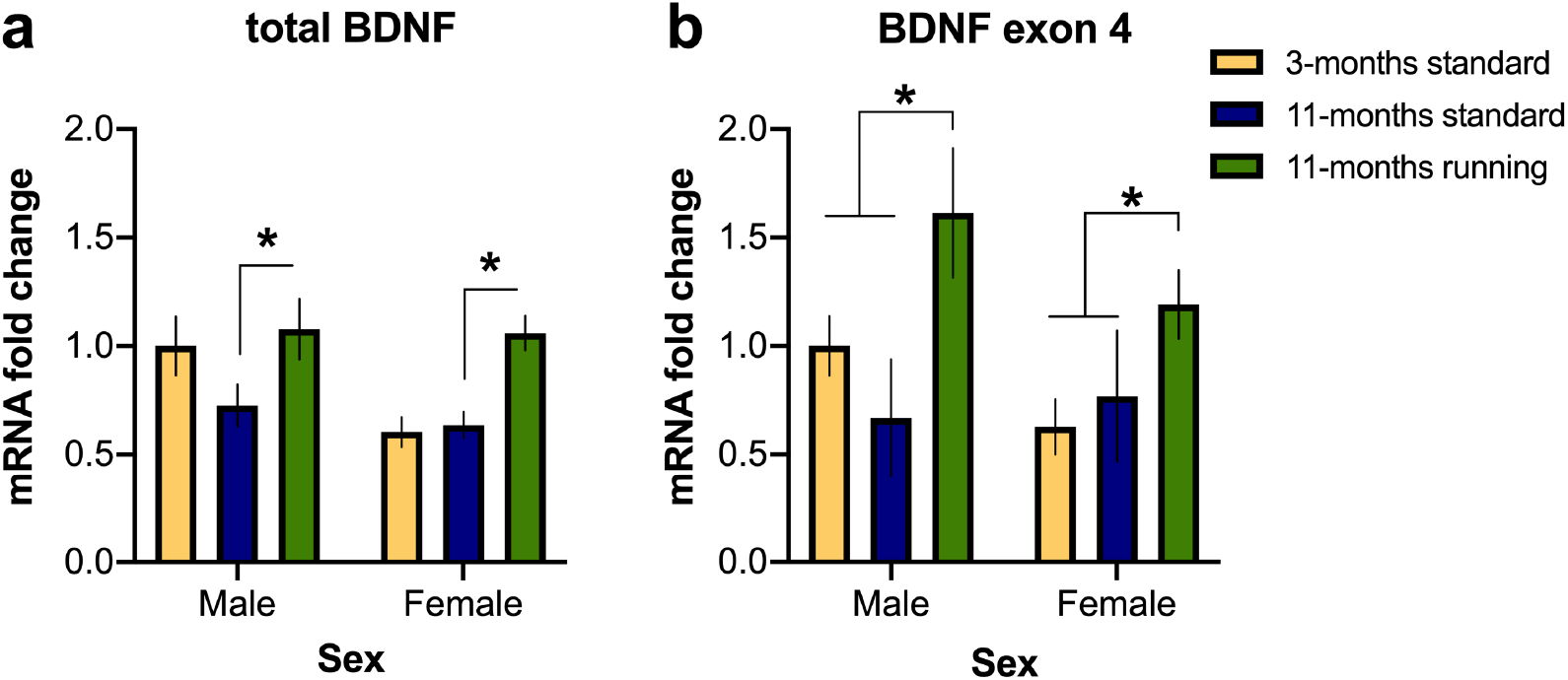
Effect of age and exercise on hippocampal *Bdnf* mRNA expression (Mean ± SEM). (a) Total *Bdnf* transcript expression in the hippocampus was significantly increased following 3 months of exercise compared to standard housing in aged mice (*p<0.05 main effect of Treatment with post-hoc Tukey comparisons). (b) *Bdnf* exon 4 transcript expression in the hippocampus was significantly increased following 3 months of exercise compared to standard housing in young and aged mice (*p<0.05 main effect of Treatment with post-hoc Tukey comparisons). n=3-4 per group.

## 4. Discussion

The main findings of the current study are 1) aging impairs reinstatement in both sexes; 2) aerobic exercise during middle age can restore reinstatement in males but not in females; and 3) exercise during middle age increases expression of total *Bdnf* and *Bdnf* exon 4 mRNA. Our secondary findings include combined effects of aging and running to increase freezing during conditioning, and aging increasing overall freezing during extinction and test. There was a subtle effect of males showing accelerated extinction acquisition compared to females, however, there were no sex effects by the final block of extinction. Notably, all the mice from the present study were obtained at 8 weeks of age, and were housed in the same facility until the end of the study, which an important detail given that most studies in aging mice were bought as retired breeders with unknown history.

### 4.1. Exercise, sex, and age effects on cognitive flexibility and hippocampal Bdnf

The present study found significant impairments in cognitive flexibility as measured by reinstatement of extinguished fear in mice at 11 months of age compared to mice at 3 months of age. A decrease in functional connectivity between the prefrontal cortex (PFC) and the hippocampus with age that may predict cognitive performance has been reported (Buckley et al., 2017). Such connectivity is hypothesized to be critical for reinstatement of extinguished fear (Ganella & Kim, 2014), hence a decreased connectivity in these extinction-related pathways would explain the present finding of age-associated impairment in fear reinstatement.

Interestingly, we found no sex effects between standard-housed groups at any age in our study. There is some evidence of sex differences within the literature regarding cognitive flexibility in young adults and even in children under the age of 3, with males generally performing better (Evans & Hampson, 2015; Overman, 2004). Consistent with this, a recent report in middle-aged marmosets showed that males are faster to acquire reversal learning (Laclair et al., 2019). These studies all highlighted the PFC as the main driver of observed sex differences in these reversal tasks, but did not undertake further investigations of the role of the hippocampus. The absence of sex differences between standard-housing groups in our study may be because reinstatement of extinguished fear relies heavily on hippocampus (Frohardt, Guarraci, & Bouton, 2000; Wilson et al., 1995), whereas the role of PFC in reinstatement is yet unknown. Importantly, while all rats given reminder froze more during test baseline than rats not given any reminder (i.e., context-shock association has been formed due to the reminder training; Figure 1C), only 3 out of 6 groups that received reminder actually showed reinstatement. This finding is consistent with previous reports that baseline freezing at test does not indicate relapse following extinction (Callaghan et al. 2011; Kim & Richardson, 2009; Park et al. 2020). Classic theories of extinction do highlight that relapse following extinction is not a simple summation between context fear and CS fear at test (Bouton & King, 1983; Bouton et al. 2002), strongly supported by recent studies showing a dissociation between context and CS-elicited freezing (Park et al. 2017b; 2020).

Strikingly, we observed that 3 months of exercise during middle age restored the age-related reinstatement deficit in males but not females. Sex-specific effects of exercise on fear extinction have been reported in young adult rats (Bouchet et al., 2017). In that study, acute exercise during extinction consolidation significantly improved extinction in males but had no effect on females (Bouchet et al., 2017). That finding is consistent with the idea that there is sex-specificity in how previous experiences impact cognitive flexibility. Studies of humans (Shields, Trainor, Lam, & Yonelinas, 2016) and mice (Laredo et al., 2015) have found that cognitive flexibility in males was more significantly impaired following a stressor, although one recent study found that female rats were more affected (Grafe, Cornfeld, Luz, Valentino, & Bhatnagar, 2017). Sex-differences in cognitive performance between men and women are also reported in pathological conditions. Women have greater cognitive decline in Alzheimer’s disease compared to males (Irvine, Laws, Gale, & Kondel, 2012), whereas in Parkinson’s disease patients, males have been described as having higher cognitive impairments than females (Nicoletti et al., 2016). There is still much to be discovered regarding sex differences in age-related cognitive decline.

In the present study, the protective effect of exercise in aging males does not appear to be driven by hippocampal *Bdnf* gene expression. While the tissue came from mice that received conditioning, extinction and test 1 week prior to tissue collection, increases in hippocampal BDNF mRNA following fear conditioning are only maintained up to 24 hours (Mizuno, Dempster, Mill, & Giese, 2012). Therefore, it is highly unlikely the mRNA findings are due to behavioral procedures, and all of the animals were treated identically. Taken together, aging did not appear to have any significant effect on the expression of *Bdnf* coding and exon 4 transcripts, but these were significantly up-regulated in both males and females following 3 months of voluntary exercise. “Including exon 4, the *Bdnf* gene contains multiple exons which undergo alternative splicing to create multiple exon-specific transcripts of *Bdnf* (Aid, Kazantseva, Piirsoo, Palm, & Timmusk, 2007; Timmusk et al., 1993). In addition to *Bdnf* exon 4 being required for the extinction of conditioned fear in rodents (Baker-Andresen et al., 2013; Bredy et al., 2007), it is also an important regulator of the activity dependent effects of BDNF protein. The exon 4 promoter contains a CRE binding site that is thought to be responsible for the cyclic adenosine monophosphate (cAMP) initiated transcription of BDNF important for experience-dependent plasticity (Ehrlich & Josselyn, 2016; Zheng & Wang, 2009). *Bdnf* plays a vital role in exercise-associated improvements of learning and memory (Cotman & Berchtold, 2002; Cotman et al., 2007; Neeper, Góauctemez-Pinilla, Choi, & Cotman, 1995; Zoladz, Pilc, & Pilc, 2010). In young adult rodents, both males (Neeper et al., 1996) and females (Berchtold et al., 2001) have increased levels of *Bdnf* mRNA within the hippocampus following voluntary wheel running.

While we describe no sex differences in *Bdnf* mRNA expression, previous work has found differences in hippocampal BDNF protein levels between males and females which differed following exercise (Venezia, Guth, Sapp, Spangenburg, & Roth, 2016). It is possible that future studies with increased power can assess more subtle effects of sex. Specifically, voluntary access to running wheels from 8 weeks of age for 20 weeks had increased hippocampal levels of mature BDNF protein in males only, however females had higher sedentary protein expression (Venezia et al., 2016). While expression of *Bdnf* has been found to decrease with age in rats (Silhol, Bonnichon, Rage, & Tapia-Arancibia, 2005) and in humans, a correlative decline with cognition has only been reported for females (Barha, Davis, et al., 2017; Komulainen et al., 2008). These indicate complex sex- and age-related interactions underlying functional BDNF expression changes related to exercise and consequential improvement in cognitive function. Future studies should consider interrogating the causal role of BDNF in cognition by manipulating its expression or receptor signaling during behavioral testing.

This study focused on the hippocampus because of its importance in the fear reinstatement/cognitive flexibility task used here following exercise in middle age in both sexes. While there was an increase in hippocampal *Bdnf* following exercise during aging in both sexes, in females there was no improvement in cognitive flexibility. This suggests that either increased *Bdnf* is not sufficient to restore age-related deficits in cognitive flexibility, or expression of hippocampal *Bdnf* is non-essential for fear reinstatement in females. Our aim of assessing females in this study has highlighted the complexity of the relationship of *Bdnf* and cognitive flexibility, but further work is required to fully understand the observed sex differences. The increases in *Bdnf* following exercise in this study suggest that there is an increase in synaptic plasticity within the hippocampus (Kuipers & Bramham, 2006). Previous work on female mice found that exercising for a longer period (6 months) than our study increased BDNF and hippocampal neurogenesis and improved spatial learning (Marlatt, Potter, Lucassen, & van Praag, 2012). Other brain regions such as the PFC, wherein dendritic spines density is increased by exercise (Brockett, LaMarca, & Gould, 2015), are also likely to contribute to improved cognitive flexibility (Gruber et al., 2010; Kim, Johnson, Cilles, & Gold, 2011).

### 4.2. Aging effects on enhanced conditioned fear expression

Across both sexes, aged animals with access to running wheels froze more overall during fear acquisition. Additionally, during extinction and test, aged animals regardless of exercise had increased fear recall. Such enhanced freezing in both the aged standard and aged exercise groups suggests an age-related enhancement of cued fear memory consolidation, while aging and exercise may also cause enhance acquisition of cued fear memory. These findings are consistent with a recent report of similar increases in freezing with cued fear in aged mice (Shoji & Miyakawa, 2019). The amygdala is a central brain region regulating the acquisition and consolidation of cued fear (Duvarci, Nader, & LeDoux, 2005; Kim et al., 2012; Orsini & Maren, 2012). While aged-related changes in amygdala structure and volume are currently poorly understood, there may be an overall loss of functional connectivity of the amygdala with the broader cortical regions (Von Bohlen und Halbach & Unsicker, 2002). However, the mechanisms whereby loss of innervation of the amygdala could result in increased conditioned fear consolidation remains to be investigated.

### 4.3. Limitations

A limitation of the study is that estrous cycle and estrogen levels of the females were not monitored. Estrous cycle has been shown to affect fear conditioning and extinction in female rodents and humans (Blume et al., 2017; Zeidan et al., 2011), with recent evidence that estrous cycle and related sex hormones drive extinction differences between male and female rats (Blume et al., 2017; Perry et al., 2020). However, the low variability in the female groups in the present study suggests that it is unlikely that estrous cycle is influencing our results. This may especially be the case for the female aging mice test at 11 months of age undergoing perimenopause with drastically reduced levels of circulating sex hormones (Diaz Brinton, 2012; Koebele & Bimonte-Nelson, 2016; Mobbs, Gee, & Finch, 1984).

In addition, running activity was not monitored, so it is unclear if the exercise levels were equivalent between males and females. Nevertheless, BDNF levels were induced to similar extents in males and females, suggesting that the running stimulus was sufficient to activate exercise-associated molecular signaling programs in the hippocampus.

### 4.4. Conclusions

Overall, our study suggests that exercise during middle age reduces the extent of cognitive impairments in males but not females. The benefits of exercise described here highlight the importance of tailoring exercise recommendations/expectations to the individual, with sex being an important consideration. Importantly, reinstatement of conditioned fear appears to be a helpful rodent model to delineate sex and age effects on cognition, and we hope that our observations will facilitate further studies to understand the molecular correlates of exercise benefits on cognitive flexibility.

## Acknowledgments

This work was supported by National Health and Medical Research Council Career Development Fellowship (1083309) and Australian Research Council Discovery Grant (150102496) awarded to JHK. AJH and TYP were supported by an NHMRC Project Grant (1083468). AJH is an NHMRC Principal Research Fellow. We have no conflicts of interest. The data of this study are available from the corresponding author upon reasonable request.

## CRediT author statement

**Annabel Short:** Conceptualization, Methodology, Formal analysis, Investigation, Writing – original draft; **Viet Bui:** Investigation, **Isabel Zbukvic:** Investigation, Writing – review & editing **Terence Pang:** Investigation, Validation, Writing – review & editing, **Anthony Hannan:** Writing – review & editing, **Jee Hyun Kim:** Conceptualization, Methodology, Formal analysis, Writing – original draft, Writing – original draft, review & editing, Supervision, Funding acquisition.

## References

Aid, T., Kazantseva, A., Piirsoo, M., Palm, K., & Timmusk, T. (2007). Mouse and rat BDNF gene structure and expression revisited. Journal of Neuroscience Research, 85, 525–535.

Anacker, C., & Hen, R. (2017, June 1). Adult hippocampal neurogenesis and cognitive flexibility-linking memory and mood. Nature Reviews Neuroscience. Nature Publishing Group.

Baker-Andresen, D., Flavell, C. R., Li, X., & Bredy, T. W. (2013). Activation of BDNF signaling prevents the return of fear in female mice. Learning & Memory, 20, 237–240.

Barha, C. K., Davis, J. C., Falck, R. S., Nagamatsu, L. S., & Liu-Ambrose, T. (2017). Sex differences in exercise efficacy to improve cognition: A systematic review and meta-analysis of randomized controlled trials in older humans. Frontiers in Neuroendocrinology, 46, 71–85.

Bell, M. R. (2018). Comparing Postnatal Development of Gonadal Hormones and Associated Social Behaviors in Rats, Mice, and Humans. Endocrinology, 159(7), 2596–2613. http://doi.org/10.1210/en.2018-00220

Berchtold, N. C., Chinn, G., Chou, M., Kesslak, J. P., & Cotman, C. W. (2005). Exercise primes a molecular memory for brain-derived neurotrophic factor protein induction in the rat hippocampus. Neuroscience, Advanced online publication. doi:10.1016/j.neuroscience.2005.03.026.

Berchtold, N. C., Kesslak, J. P., Pike, C. J., Adlard, P. A., & Cotman, C. W. (2001). Estrogen and exercise interact to regulate brain-derived neurotrophic factor mRNA and protein expression in the hippocampus. European Journal of Neuroscience, 14, 1992–2002.

Blume, S. R., Freedberg, M., Vantrease, J. E., Chan, R., Padival, M., Record, M. J., et al. (2017). Sex- and estrus-dependent differences in rat basolateral amygdala. Journal of Neuroscience, 0758–17–20. http://doi.org/10.1523/JNEUROSCI.0758-17.2017

Bouchet, C. A., Lloyd, B. A., Loetz, E. C., Farmer, C. E., Ostrovskyy, M., Haddad, N., … Greenwood, B. N. (2017). Acute exercise enhances the consolidation of fear extinction memory and reduces conditioned fear relapse in a sex-dependent manner. Learning and Memory, 24, 358–368.

Bouton, M. E. (2002). Context, ambiguity, and unlearning: sources of relapse after behavioral extinction. Biological Psychiatry, 52(10), 976–986.

Bouton, M. E., & King, D. A. (1983). Contextual control of the extinction of conditioned fear: Tests for the associative value of the context. Journal of Experimental Psychology: Animal Behavior Processes, 9(3), 248–265. http://doi.org/10.1037//0097-7403.9.3.248

Bredy, T. W., Wu, H., Crego, C., Zellhoefer, J., Sun, Y. E., & Barad, M. (2007). Histone modifications around individual BDNF gene promoters in prefrontal cortex are associated with extinction of conditioned fear. Learning & Memory, 14, 268–276.

Brockett, A. T., LaMarca, E. A., & Gould, E. (2015). Physical Exercise Enhances Cognitive Flexibility as Well as Astrocytic and Synaptic Markers in the Medial Prefrontal Cortex. PLOS ONE, 10, e0124859.

Buckley, R. F., Schultz, A. P., Hedden, T., Papp, K. V., Hanseeuw, B. J., Marshall, G., … Chhatwal, J. P. (2017). Functional network integrity presages cognitive decline in preclinical Alzheimer disease. Neurology, 89, 29–37.

Burghardt, N. S., Park, E. H., Hen, R., & Fenton, A. A. (2012). Adult-born hippocampal neurons promote cognitive flexibility in mice. Hippocampus, 22, 1795–1808.

Callaghan, B. L., & Richardson, R. (2011). Maternal separation results in early emergence of adult-like fear and extinction learning in infant rats. Behavioral Neuroscience, 125(1), 20–28. http://doi.org/10.1037/a0022008

Chaddock, L., Erickson, K. I., Prakash, R. S., Kim, J. S., Voss, M. W., Vanpatter, M., … Kramer, A. F. (2010). A neuroimaging investigation of the association between aerobic fitness, hippocampal volume, and memory performance in preadolescent children. Brain Research, Advanced online publication. doi:10.1016/j.brainres.2010.08.049.

Chen, N. A., Ganella, D. E., Bathgate, R. A. D., Chen, A., Lawrence, A. J., & Kim, J. H. (2016). Knockdown of corticotropin-releasing factor 1 receptors in the ventral tegmental area enhances conditioned fear. European Neuropsychopharmacology: the Journal of the European College of Neuropsychopharmacology, 26(9), 1533–1540. http://doi.org/10.1016/j.euroneuro.2016.06.002

Cotman, C. W., & Berchtold, N. C. (2002, June 1). Exercise: A behavioral intervention to enhance brain health and plasticity. Trends in Neurosciences.

Cotman, C. W., Berchtold, N. C., & Christie, L.-A. A. (2007). Exercise builds brain health: key roles of growth factor cascades and inflammation. Trends in Neurosciences, 30, 464–472.

Dajani, D. R., & Uddin, L. Q. (2015). Demystifying cognitive flexibility: Implications for clinical and developmental neuroscience. Trends in Neurosciences, 38, 571–578.

Diaz Brinton, R. (2012). Minireview: translational animal models of human menopause: challenges and emerging opportunities. Endocrinology, 153(8), 3571–3578. http://doi.org/10.1210/en.2012-1340

Duvarci, S., Nader, K., & LeDoux, J. E. (2005). Activation of extracellular signal-regulated kinase-mitogen-activated protein kinase cascade in the amygdala is required for memory reconsolidation of auditory fear conditioning. The European Journal of Neuroscience, 21, 283–289.

Ehrlich, D. E., & Josselyn, S. A. (2016). Plasticity-related genes in brain development and amygdala-dependent learning. Genes, Brain and Behavior, 15, 125–143.

Erickson, K. I., Voss, M. W., Prakash, R. S., Basak, C., Szabo, A., Chaddock, L., … Kramer, A. F. (2011). Exercise training increases size of hippocampus and improves memory. Proceedings of the National Academy of Sciences of the United States of America, Advanced online publication. doi:10.1073/pnas.1015950108.

Evans, K. L., & Hampson, E. (2015). Sex differences on prefrontally-dependent cognitive tasks. Brain and Cognition, Advanced online publication. doi:10.1016/j.bandc.2014.11.006.

Falls, W. A., Fox, J. H., & MacAulay, C. M. (2010). Voluntary exercise improves both learning and consolidation of cued conditioned fear in C57 mice. Behavioural Brain Research, 207(2), 321–331. http://doi.org/10.1016/j.bbr.2009.10.016

Ferreira, L., Ferreira Santos-Galduróz, R., Ferri, C. P., & Fernandes Galduróz, J. C. (2014). Rate of cognitive decline in relation to sex after 60 years-of-age: A systematic review. Geriatrics & Gerontology International, 14, 23–31.

Frohardt, R. J., Guarraci, F. A., & Bouton, M. E. (2000). The effects of neurotoxic hippocampal lesions on two effects of context after fear extinction. Behavioral Neuroscience, 114, 227–240.

Ganella, D. E., & Kim, J. H. (2014). Developmental rodent models of fear and anxiety: from neurobiology to pharmacology. British Journal of Pharmacology, 171, 4556–4574.

Giovanello, K. S., Schnyer, D., & Verfaellie, M. (2009). Distinct hippocampal regions make unique contributions to relational memory. Hippocampus, 19(2), 111–117. http://doi.org/10.1002/hipo.20491

Grafe, L. A., Cornfeld, A., Luz, S., Valentino, R., & Bhatnagar, S. (2017). Orexins Mediate Sex Differences in the Stress Response and in Cognitive Flexibility. Biological Psychiatry, 81, 683–692.

Gruber, A. J., Calhoon, G. G., Shusterman, I., Schoenbaum, G., Roesch, M. R., & O’Donnell, P. (2010). More is less: A disinhibited prefrontal cortex impairs cognitive flexibility. Journal of Neuroscience, 30, 17102–17110.

Handford, C. E., Tan, S., Lawrence, A. J., & Kim, J. H. (2014). The effect of the mGlu5 negative allosteric modulator MTEP and NMDA receptor partial agonist D-cycloserine on Pavlovian conditioned fear. The International Journal of Neuropsychopharmacology / Official Scientific Journal of the Collegium Internationale Neuropsychopharmacologicum (CINP), 17, 1521–1532.

Herting, M. M., & Nagel, B. J. (2012). Aerobic fitness relates to learning on a virtual Morris Water Task and hippocampal volume in adolescents. Behavioural Brain Research, Advanced online publication. doi:10.1016/j.bbr.2012.05.012.

Holland, P. C., & Bouton, M. E. (1999). Hippocampus and context in classical conditioning. Current Opinion in Neurobiology, 9, 195–202.

Irvine, K., Laws, K. R., Gale, T. M., & Kondel, T. K. (2012). Greater cognitive deterioration in women than men with Alzheimer’s disease: A meta analysis. Journal of Clinical and Experimental Neuropsychology, 34, 989–998.

Jonasson, Z. (2005). Meta-analysis of sex differences in rodent models of learning and memory: a review of behavioral and biological data. Neuroscience & Biobehavioral Reviews, 28, 811–825.

Karlsson, P., Thorvaldsson, V., Skoog, I., Gudmundsson, P., & Johansson, B. (2015). Birth cohort differences in fluid cognition in old age: Comparisons of trends in levels and change trajectories over 30 years in three population-based samples. Psychology and Aging, 30, 83–94.

Kim, C., Johnson, N. F., Cilles, S. E., & Gold, B. T. (2011). Common and distinct mechanisms of cognitive flexibility in prefrontal cortex. Journal of Neuroscience, 31, 4771–4779.

Kim, J. H., Li, S., Hamlin, A. S., McNally, G. P., & Richardson, R. (2012). Phosphorylation of mitogen-activated protein kinase in the medial prefrontal cortex and the amygdala following memory retrieval or forgetting in developing rats. Neurobiology of Learning and Memory, 97(1), 59–68. http://doi.org/10.1016/j.nlm.2011.09.005

Kim, J. H., & Richardson, R. (2007). A developmental dissociation in reinstatement of an extinguished fear response in rats. Neurobiology of Learning and Memory, 88, 48–57.

Kim, J. H., & Richardson, R. (2009). Expression of renewal is dependent on the extinction-test interval rather than the acquisition-extinction interval. Behavioral Neuroscience, 123(3), 641–649. http://doi.org/10.1037/a0015237

Kim, J. H., & Richardson, R. (2010). New findings on extinction of conditioned fear early in development: theoretical and clinical implications. Biological Psychiatry, 67, 297–303.

Kleemeyer, M. M., Kühn, S., Prindle, J., Bodammer, N. C., Brechtel, L., Garthe, A., … Lindenberger, U. (2016). Changes in fitness are associated with changes in hippocampal microstructure and hippocampal volume among older adults. NeuroImage, Advanced online publication. doi:10.1016/j.neuroimage.2015.11.026.

Koebele, S. V., & Bimonte-Nelson, H. A. (2016). Modeling menopause: The utility of rodents in translational behavioral endocrinology research. Maturitas, 87, 5–17. http://doi.org/10.1016/j.maturitas.2016.01.015

Komulainen, P., Pedersen, M., Hänninen, T., Bruunsgaard, H., Lakka, T. A., Kivipelto, M., … Rauramaa, R. (2008). BDNF is a novel marker of cognitive function in ageing women: The DR’s EXTRA Study. Neurobiology of Learning and Memory, 90, 596–603.

Kuipers, S. D., & Bramham, C. R. (2006). Brain-derived neurotrophic factor mechanisms and function in adult synaptic plasticity: new insights and implications for therapy. Current Opinion in Drug Discovery & Development, 9, 580–586.

Laclair, M., Febo, M., Nephew, B., Gervais, N. J., Poirier, G., Workman, K., … Lacreuse, A. (2019). Cognition and Behavior Sex Differences in Cognitive Flexibility and Resting Brain Networks in Middle-Aged Marmosets, Advanced online publication. doi:10.1523/ENEURO.0154-19.2019.

Laredo, S. A., Steinman, M. Q., Robles, C. F., Ferrer, E., Ragen, B. J., & Trainor, B. C. (2015). Effects of defeat stress on behavioral flexibility in males and females: modulation by the mu-opioid receptor. European Journal of Neuroscience, 41, 434–441.

Linn, M. C., & Petersen, A. C. (1985). Emergence and Characterization of Sex Differences in Spatial Ability: A Meta-Analysis. Child Development, 56, 1479.

Lynch, G., Rex, C. S., & Gall, C. M. (2006). Synaptic plasticity in early aging. Ageing Research Reviews, 5(3), 255–280. http://doi.org/10.1016/j.arr.2006.03.008

Madsen, H. B., Guerin, A. A., & Kim, J. H. (2017). Investigating the role of dopamine receptor- and parvalbumin-expressing cells in extinction of conditioned fear. Neurobiology of Learning and Memory, 145, 7–17.

Madsen, H. B., & Kim, J. H. (2016). Ontogeny of memory: An update on 40 years of work on infantile amnesia. Behavioural Brain Research, 298(Part A), 4–14. http://doi.org/10.1016/j.bbr.2015.07.030

Maren, S., Phan, K. L., & Liberzon, I. (2013). The contextual brain: Implications for fear conditioning, extinction and psychopathology. Nature Reviews. Neuroscience, 14(6), 417–428. http://doi.org/10.1038/nrn3492

Marlatt, M. W., Potter, M. C., Lucassen, P. J., & van Praag, H. (2012). Running throughout middle-age improves memory function, hippocampal neurgenesis and BDNF levels in female C57Bl/6J mice. Developmental Neurobiology, 72, 943–952.

Matsuda, S., Matsuzawa, D., Ishii, D., Tomizawa, H., Sutoh, C., & Shimizu, E. (2015). Sex differences in fear extinction and involvements of extracellular signal-regulated kinase (ERK), Advanced online publication. doi:10.1016/j.nlm.2015.05.009.

McCarrey, A. C., An, Y., Kitner-Triolo, M. H., Ferrucci, L., & Resnick, S. M. (2016). Sex differences in cognitive trajectories in clinically normal older adults. Psychology and Aging, 31, 166–175.

Mizuno, K., Dempster, E., Mill, J., & Giese, K. P. (2012). Long-lasting regulation of hippocampal Bdnf gene transcription after contextual fear conditioning. Genes, Brain and Behavior, 11(6), 651–659. http://doi.org/10.1111/j.1601-183X.2012.00805.x

Mobbs, C. V., Gee, D. M., & Finch, C. E. (1984). Reproductive senescence in female C57BL/6J mice: ovarian impairments and neuroendocrine impairments that are partially reversible and delayable by ovariectomy. Endocrinology, 115(5), 1653–1662. http://doi.org/10.1210/endo-115-5-1653

Neeper, S. A., Góauctemez-Pinilla, F., Choi, J., & Cotman, C. (1995). Exercise and brain neurotrophins. Nature, 373, 109–109.

Neeper, S. A., Gómez-Pinilla, F., Choi, J., & Cotman, C. W. (1996). Physical activity increases mRNA for brain-derived neurotrophic factor and nerve growth factor in rat brain. Brain Research, 726, 49–56.

Nicoletti, A., Vasta, R., Mostile, G., Nicoletti, G., Arabia, G., Iliceto, G., … Zappia, M. (2016). Gender effect on non-motor symptoms in Parkinson’s disease: are men more at risk?, Advanced online publication. doi:10.1016/j.parkreldis.2016.12.008.

Nishijima, T., Llorens-Martín, M., Tejeda, G. S., Inoue, K., Yamamura, Y., Soya, H., et al. (2013). Cessation of voluntary wheel running increases anxiety-like behavior and impairs adult hippocampal neurogenesis in mice. Behavioural Brain Research, 245, 34–41. http://doi.org/10.1016/j.bbr.2013.02.009

Orsini, C. a, & Maren, S. (2012). Neural and cellular mechanisms of fear and extinction memory formation. Neuroscience and Biobehavioral Reviews, 36, 1773–1802.

Overman, W. H. (2004). Sex differences in early childhood, adolescence, and adulthood on cognitive tasks that rely on orbital prefrontal cortex. Brain and Cognition, 55, 134–147.

Park, C. H. J., Ganella, D. E., & Kim, J. H. (2017). Juvenile female rats, but not male rats, show renewal, reinstatement, and spontaneous recovery following extinction of conditioned fear. Learning and Memory, Advanced online publication. doi:10.1101/lm.045831.117.

Park, C. H. J., Ganella, D. E., & Kim, J. H. (2017). A dissociation between renewal and contextual fear conditioning in juvenile rats. Developmental Psychobiology, 59(4), 515–522. http://doi.org/10.1002/dev.21516

Park, C. H. J., Ganella, D. E., & Kim, J. H. (2020a). Context fear learning and renewal of extinguished fear are dissociated in juvenile female rats. Developmental Psychobiology, 62(1), 123–129. http://doi.org/10.1002/dev.21888

Perry, C. J., Ganella, D. E., Nguyen, L. D., Du, X., Drummond, K. D., Whittle, S., et al. (2020). Assessment of conditioned fear extinction in male and female adolescent rats. Psychoneuroendocrinology, in press.

Rosano, C., Guralnik, J., Pahor, M., Glynn, N. W., Newman, A. B., Ibrahim, T. S., … Aizenstein, H. J. (2017). Hippocampal Response to a 24-Month Physical Activity Intervention in Sedentary Older Adults. American Journal of Geriatric Psychiatry, Advanced online publication. doi:10.1016/j.jagp.2016.11.007.

Rubin, R. D., Watson, P. D., Duff, M. C., & Cohen, N. J. (2014). The role of the hippocampus in flexible cognition and social behavior. Frontiers in Human Neuroscience.

Salthouse, T. A. (1996). The processing-speed theory of adult age differences in cognition. Psychological Review, 103, 403–428.

Scott, W. A. (1962). Cognitive complexity and cognitive flexibility. Sociometry, 25, 405–414.

Shields, G. S., Trainor, B. C., am, J. C. W., & Yonelinas, A. P. (2016). Acute stress impairs cognitive flexibility in men, not women. Stress, 19, 542–546.

Shoji, H., & Miyakawa, T. (2019). Age-related behavioral changes from young to old age in male mice of a C57BL/6J strain maintained under a genetic stability program. Neuropsychopharmacology Reports, 39, 100–118.

Short, A. K., Fennell, K. A., Perreau, V. M., Fox, A., O’Bryan, M. K., Kim, J. H., … Hannan, A. J. (2016). Elevated paternal glucocorticoid exposure alters the small noncoding RNA profile in sperm and modifies anxiety and depressive phenotypes in the offspring. Translational Psychiatry, 6, e837.

Short, A. K., Yeshurun, S., Powell, R., Perreau, V. M., Fox, A., Kim, J. H., … Hannan, A. J. (2017). Exercise alters mouse sperm small noncoding RNAs and induces a transgenerational modification of male offspring conditioned fear and anxiety. Translational Psychiatry, 7, e1114.

Silhol, M., Bonnichon, V., Rage, F., & Tapia-Arancibia, L. (2005). Age-related changes in brain-derived neurotrophic factor and tyrosine kinase receptor isoforms in the hippocampus and hypothalamus in male rats. Neuroscience, 132, 613–624.

Stillman, C. M., Uyar, F., Huang, H., Grove, G. A., Watt, J. C., Wollam, M. E., & Erickson, K. I. (2018). Cardiorespiratory fitness is associated with enhanced hippocampal functional connectivity in healthy young adults. Hippocampus, Advanced online publication. doi:10.1002/hipo.22827.

Thomas, A. G., Dennis, A., Rawlings, N. B., Stagg, C. J., Matthews, L., Morris, M., … Johansen-Berg, H. (2016). Multi-modal characterization of rapid anterior hippocampal volume increase associated with aerobic exercise. NeuroImage, Advanced online publication. doi:10.1016/j.neuroimage.2015.10.090.

Timmusk, T., Palm, K., Metsis, M., Reintam, T., Paalme, V., Saarma, M., & Persson, H. (1993). Multiple promoters direct tissue-specific expression of the rat BDNF gene. Neuron, 10, 475–489.

van Praag, H., Shubert, T., Zhao, C., & Gage, F. H. (2005). Exercise enhances learning and hippocampal neurogenesis in aged mice. The Journal of Neuroscience: The Official Journal of the Society for Neuroscience, 25, 8680–8685.

Venezia, A. C., Guth, L. M., Sapp, R. M., Spangenburg, E. E., & Roth, S. M. (2016). Sex-dependent and independent effects of long-term voluntary wheel running on Bdnf mRNA and protein expression. Physiology and Behavior, 156, 8–15.

Von Bohlen und Halbach, O., & Unsicker, K. (2002). Morphological alterations in the amygdala and hippocampus of mice during ageing. European Journal of Neuroscience, 16, 2434–2440.

Voulo, M. E., & Parsons, R. G. (2017). Response-specific sex difference in the retention of fear extinction. Learning & Memory, 24, 245–251.

Voyer, D., Voyer, S., & Bryden, M. P. (1995). Magnitude of sex differences in spatial abilities: A meta-analysis and consideration of critical variables. Psychological Bulletin, 117, 250–270.

Wilson, A., Brooks, D. C., & Bouton, M. E. (1995). The role of the rat hippocampal system in several effects of context in extinction. Behavioral Neuroscience, 109, 828–836.

Workman, K. P., Healey, B., Carlotto, A., & Lacreuse, A. (2019). One-year change in cognitive flexibility and fine motor function in middle-aged male and female marmosets (*Callithrix jacchus*). American Journal of Primatology, 81, e22924.

Yagi, S., & Galea, L. A. M. (2019). Sex differences in hippocampal cognition and neurogenesis. Neuropsychopharmacology, 44, 200–213.

Zaninotto, P., Batty, G. D., Allerhand, M., & Deary, I. J. (2018). Cognitive function trajectories and their determinants in older people: 8 Years of follow-up in the English Longitudinal Study of Ageing. Journal of Epidemiology and Community Health, 72, 685–694.

Zanos, P., Bhat, S., Terrillion, C. E., Smith, R. J., Tonelli, L. H., & Gould, T. D. (2015). Sex-dependent modulation of age-related cognitive decline by the L-type calcium channel gene Cacna1c (Ca v 1.2). European Journal of Neuroscience, 42, 2499–2507.

Zeidan, M. A., Igoe, S. A., Linnman, C., Vitalo, A., Levine, J. B., Klibanski, A., et al. (2011). Estradiol Modulates Medial Prefrontal Cortex and Amygdala Activity During Fear Extinction in Women and Female Rats. Biological Psychiatry, 70(10), 920–927. http://doi.org/10.1016/j.biopsych.2011.05.016

Zheng, F., & Wang, H. (2009). NMDA-mediated and self-induced bdnf exon IV transcriptions are differentially regulated in cultured cortical neurons. Neurochemistry International, 54, 385–392.

Zoladz, J. A., Pilc, A., & Pilc, J. (2010). The effect of physical activity on the brain derived neurotrophic factor: from animal to human studies. Journal of Physiology and Pharmacolology, 61, 533–541.

